# Top-down perceptual modulation of pattern but not component tuned motion circuits in V1

**DOI:** 10.1101/2023.12.16.571695

**Authors:** Kim Beneyton, Micha Heilbron, Henry Kennedy, Kenneth Knoblauch, Floris P. de Lange

## Abstract

Top-down influences play a central role in perception. In multi-stable sensory phenomena subjective interpretation fluctuates spontaneously over time and the perceptual switches reflect internally driven representations engaging perceptual decision-making. Using functional magnetic resonance imaging (fMRI), we investigated how perceptual states are represented across early visual areas during perception of a bistable moving plaid. While subjects maintained central fixation, perception of the stimulus—two superimposed, obliquely oriented moving gratings—spontaneously alternated between component motion (2 gratings sliding obliquely over each other) and pattern motion (a single coherent pattern moving laterally). Direction-selective clusters were reliably modulated by perceptual state across both early (V1) and higher-order (V2 and the human medial temporal complex (hMT+)) visual regions. Pattern modulation was significantly stronger in primary visual cortex (V1) than in higher-order areas, suggesting that coherent percept formation engages mechanisms beyond simple bottom-up feature integration. Enhanced V1 modulation points to a prominent role for top-down influences from higher-order regions, consistent with predictive coding frameworks in which higher-level areas convey predictions to lower levels to disambiguate sensory input and drive perceptual inference under ambiguous conditions. Eye tracking confirmed that reflexive eye movement direction did not predict perceptual state, excluding an oculomotor confound. Interpretation of the results in light of current knowledge of the physiology of motion integration in early visual areas suggests circuits at which feedback would be predicted to act.

## Introduction

Sensory input is inherently ambiguous: a given pattern of sensory stimulation can be consistent with multiple interpretations. In vision, a classic example of this ambiguity arises in motion processing, where the relationship between spatiotemporal variations on the retina and perceived motion direction and speed is underdetermined at the level of local motion detectors [1–3]. In primary visual cortex, area V1, the trade-offs between the direction and speed of drifting gratings can lead to the same responses to gratings moving in different directions and at different speeds in univariant, direction-selective cells [4]. It is the comparison of direction-specific responses over an ensemble of such differentially tuned cells, however, that provides the information necessary to disambiguate motion speed and direction. Such comparisons are thought to be performed at higher-order levels in the visual cortex. In this manner, the ambiguity in responses of cells in early visual areas has been proposed to be resolved by higher-order visual motion areas, including the Medial Temporal (MT/V5) [5], Medial Superior Temporal (MST) areas [6] and the medial motion areas (areas V6 and V6A) [7].

Evidence for hierarchical, bottom-up processing comes from studies investigating the perception of moving plaid stimuli in human and non-human primates. Superimposed drifting gratings of different orientation cohere to a plaid pattern that moves in a direction different from that of either component grating alone [2]. Early electrophysiological studies using such stimuli in anesthetized animals reported that cells in visual areas V1 and V2 exhibited direction-tuning curves that reflected the component directions, whereas increasing numbers of cells in both areas MT and MST responded selectively to the direction of the pattern motion [4]. Accordingly, it was expected that the representation of pattern motion is limited to higher-order visual areas such as MT and MST while it was thought to be absent from area V1. Subsequent studies however, in awake and behaving animals, reported small numbers of pattern motion-selective cells as early as area V1 [8–10]. More recently, single-cell responses to complex pattern features have also been obtained using large-scale two-photon cortical imaging in the upper layers of awake macaque V1 [11]. In particular, Guan et al. [12] reported a large proportion of plaid orientation-selective detectors in macaque area V1 of which ∼5% exhibited direction-selectivity.

In humans, multiple fMRI studies that investigated pattern motion responses across the visual cortical hierarchy (from V1 to hMT+) reported robust evidence for pattern-selective responses in extrastriate regions, such as V3 and hMT+ [13, 14]; by contrast, pattern selectivity in early visual areas like V1 and V2 was weak or undetected. These findings were consistent with the hierarchical framework in which motion integration becomes progressively more robust at successive stages of the dorsal visual pathway. However, it is important to note that methodological choices can strongly influence what motion signals are detectable. For example, adaptation paradigms such as that used in [13] are effective at isolating strong pattern motion responses in hMT+ but may lack sensitivity for detecting relatively small and spatially distributed signals in V1 and V2. Similarly, without an independent direction-selective localizer to define selective clusters, the univariate analyses used in [14] were too coarse to reveal fine scale motion direction selectivity.

Recently, multivariate pattern analysis across the visual cortical hierarchy was successfully exploited to decode the perceived direction of moving plaid stimuli as early as area V1 [15]. In a subsequent study using an occlusion paradigm [16], it was further demonstrated that percept-related signals were decodable from non-stimulated regions of early visual cortex, findings the authors interpreted as consistent with top-down influences. However, because the stimulus differed between component and pattern conditions and the decoded directional information overlapped, bottom-up contributions could not be entirely excluded. The authors also acknowledged that attentional modulation might have driven percept related effects in non-stimulated regions, leaving the precise source of decodable signals ambiguous.

Recent high resolution fMRI studies using independent direction-selective localizers have revealed motion direction selective clusters in the context of ambiguous motion, particularly within hMT+ (e.g., [17]), and there is converging evidence that early visual cortex, including V1, reflects top-down influences during perceptual ambiguity [18, 19]. The discrepancy between early and recent studies highlights the importance of two key methodological factors in detecting pattern motion selectivity in early visual areas: (1) independently identifying direction-tuned voxel clusters to capture fine-grained percept-related signals, and (2) maintaining a constant bottom-up sensory input across conditions in ambiguous motion paradigms to isolate perceptual effects from sensory differences.

Given that both electrophysiological and fMRI findings converge on the presence of pattern-motion signals as early as area V1, an important question is: what is the origin of these signals? In non-human primates, individual neurons in area V1 have relatively small receptive fields and the intrinsic connections within V1 extend over limited spatial extents [20], which makes it unlikely that the large scale integration required for pattern motion selectivity is intrinsic to area V1. Neurocomputational modeling has contributed to clarify how interactions between different types of V1 inputs could bias MT neurons toward either component or pattern motion responses. For example, models that combine standard direction-selective V1 cells, end-stopped V1 neurons, and V1 cells with suppressive extra-classical receptive fields demonstrate that varying the relative strength of these inputs can shift MT responses along the continuum from component-like to pattern-like selectivity [21]. In these frameworks, end-stopped inputs—which are sensitive to terminators such as line endings and corners—can provide locally unambiguous motion cues that are useful for motion integration in MT. Nevertheless, while V1 inputs may help sharpen direction tuning in MT, they cannot explain the spontaneous alternations between competing percepts seen during bistable motion perception. Instead, these spontaneous perceptual switches are generally thought to arise from competitive interactions among neural populations representing alternative interpretations of ambiguous input, involving dynamic interactions across multiple processing levels and top-down influences [22, 23].

The top-down projections from visual motion areas including area MT to V1 are surprisingly strong, given the large anatomical [24] and hierarchical [25, 26] distances that separate them. Therefore, a possibility is that pattern responses in area V1 are not computed locally but derived from feedback from MT and higher-order areas [25, 27, 28] and would align with recent high-resolution studies identifying a role of top-down signals in inducing internally-driven mental images [29, 30].

In the current study, we sought to isolate internally-generated signals from bottom-up induced influences in order to investigate changes in brain states across the cortical hierarchy during spontaneous perceptual alternations of a moving plaid stimulus. A bistable motion paradigm is ideal for studying brain state changes due to intrinsic processing since the external driving stimulation parameters remain fixed while the percept shifts between alternative interpretations [23]. To disentangle the effects of the physical stimulus from that of the perceptual interpretation, we compared activity during intervals in which subjects reported pattern or component percepts in three hierarchically distinct visual areas, using a moving plaid. Direction-specific functional localizers were used to determine voxels selective for a particular motion direction in areas V1, V2 and hMT+ that were subsequently used as regions of interest to estimate blood oxygen level dependent (BOLD) contrasts between the pattern and the component perceptual states. To preview, the results show that voxel clusters in all three regions, including V1, reliably tracked the current perceptual state, in support of an important role of top-down signals in constructing perceptual experience. Interestingly, the regions modulated were specific to a particular perceptual state in each cortical area, with only the pattern-selective voxels modulated within V1.

## Results

During the test stimulus presentation, observers’ perception randomly alternated between component and pattern motion directions (Fig 1A), analogous to the systematic alternations seen in the localizer sequence (Figure A in S1 Text). We aimed to determine whether the activity in direction-selective clusters that primarily responded to physical motion would correlate with the motion direction during the two distinct perceptual states of the bistable stimulus. To ensure that the BOLD signal could adequately capture percept-specific responses, we first compared the durations of the component and pattern states to evaluate whether the induced percept duration allowed event-related fMRI analysis of perceptual switching behavior.

**Fig 1.**
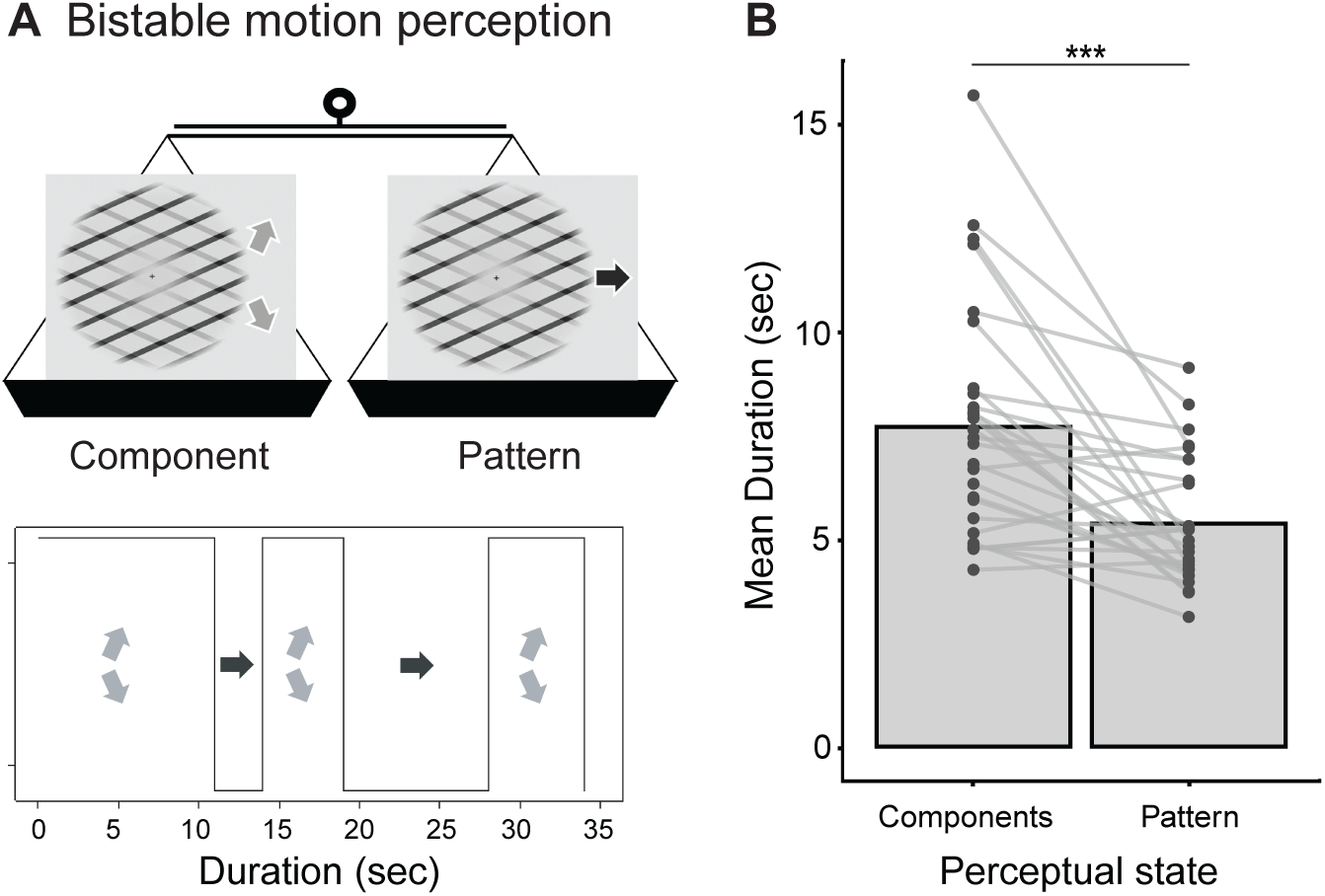
Experimental design. A. Schema of the main experimental paradigm. A bistable moving plaid was presented over an extended period of time (60 s); perceptual switching spontaneously occurred between two competing perceptual states over time: observers reported seeing either a plaid moving rightward (“pattern” percept) or two distinct component gratings moving obliquely (“component” percept). B. Group average durations for each perceptual state. Each dot represents one individual, with lines connecting within-subject paired data across conditions. Three stars (in this and other figures) indicate significant differences at *p* < 0.001 as described in the text.

Fig 1B shows the average duration of component and pattern percepts for each subject. The group average durations revealed a significantly shorter time for the pattern percept than for component percept (*t*(27) = −4.71, *p* < .001). Group average durations and their confidence intervals (pattern: mean [CI95] = 5.39 [4.84; 5.95], component: mean [CI95] = 7.73 [6.67; 8.79]) exceeded the ∼4 s minimum considered adequate for event-related BOLD estimation [31, 32].

### Differential BOLD activity for pattern/component perceptual states

To test whether the perceptual states defined by the subjective reports were accompanied by different BOLD signals in early visual areas, we selected oblique- and rightward-selective clusters within visual areas V1, V2 and hMT+ (S1 Text, S2 Text and S3 Text) and estimated the GLM contrast of BOLD signals during pattern and component states within these regions of interest (ROI). Specifically, we asked whether the pattern minus component selectivity *z*-score differed across direction-selective units (Table B in S3 Text). The statistical significance of this contrast was tested by comparing the goodness of fit of two nested linear mixed-effects (LME) models, with and without the region-based perceptual effect, using a likelihood ratio test (LRT), including all three visual areas. We found a highly significant interaction between perceptual state and direction selectivity when comparing rightward- and oblique-selective voxels (LRT (all areas): *χ*^2^ = 36.47, *df* = 3, *p* < 0.001) (Fig 2), indicating that perceptual state modulates BOLD responses in a direction-selective manner. Post hoc pattern-minus-component contrasts showed that rightward-selective voxels exhibited greater responses than oblique-selective voxels in V1 (*z* = 3.57, *p* = 0.002), V2 (*z* = 3.64, *p* = 0.001), and hMT+ (*z* = 3.78, *p* < 0.001). Oblique-selective clusters in V2 and hMT+ exhibited a significant preference for component motion, as indicated by negative z values (V2: *z* = −2.73, *p* = 0.036; hMT+: *z* = −4.02, *p* < 0.001), whereas no such preference was observed in V1 (*z* = −1.15, *p* = 0.795). In contrast, rightward-selective clusters showed enhanced activation during the pattern motion state in V1 (*z* = 3.17, *p* = 0.009), but not in V2 (*z* = 1.68, *p* = 0.412) or in hMT+ (*z* = 0.59, *p* = 0.990). These results indicate a systematic interaction between direction selectivity and perceptual state across early visual cortex.

**Fig 2.**
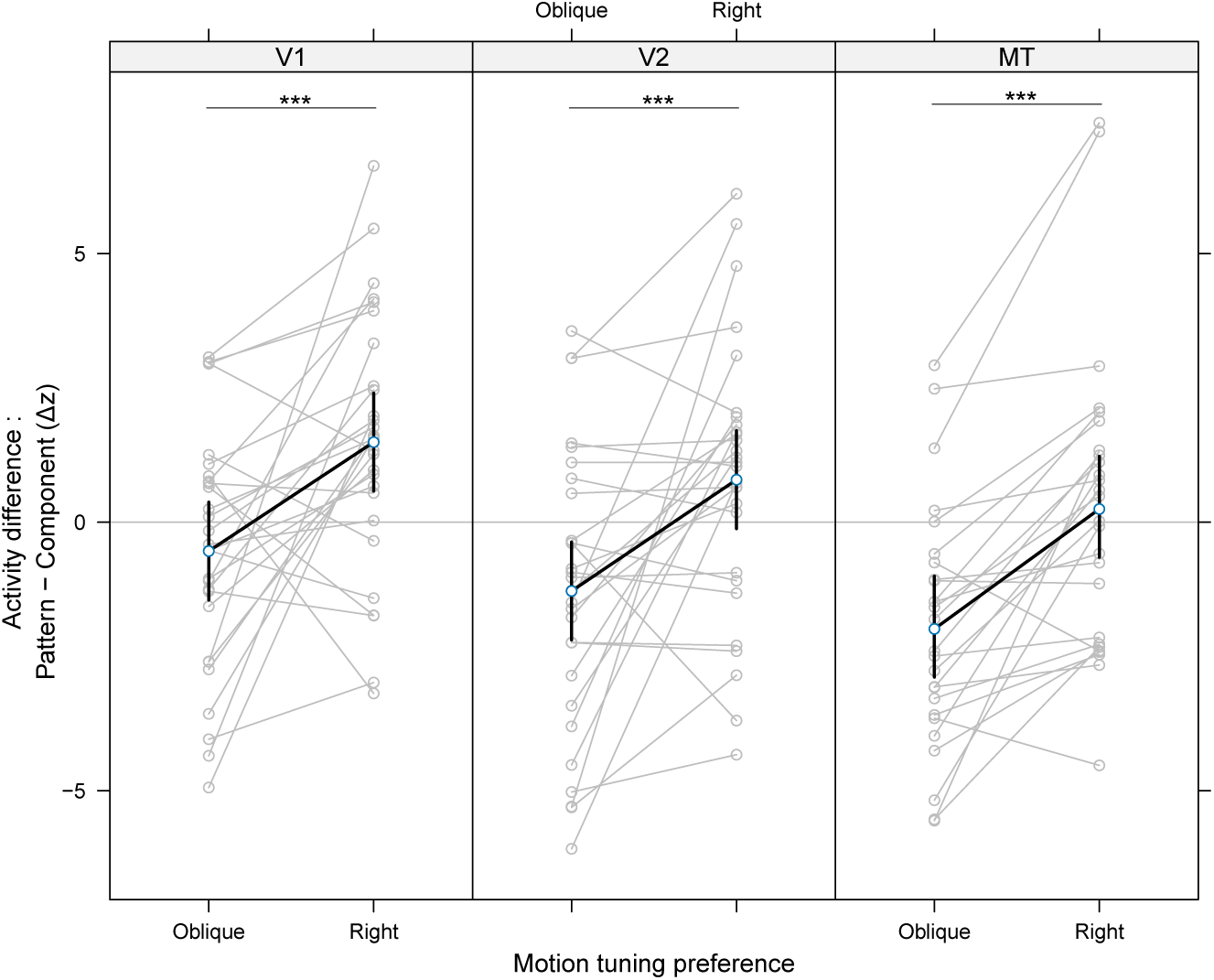
Direction-selective regions reflect perceptual state. Model estimates of contrast (pattern minus component perceptual states) within rightward and oblique direction-selective clusters in V1, V2 and hMT+ at the group-level (N=28 observers). Grey connected points indicate individual observers; black points and lines show fixed-effect estimates with 95% confidence intervals.

### Event related temporal dynamics aligned to perceptual switching

For the localizer, the difference in event-related averages (ERA) between rightward and oblique trials is plotted over time for the direction-selective clusters in V1, V2, and hMT+ (Fig 3A). In both V1 and V2, the rightward-minus-oblique differential activity varied systematically across clusters: rightward motion elicited stronger responses in rightward-selective clusters (solid lines), whereas oblique motion elicited stronger responses in oblique-selective clusters (dashed lines). This divergence between cluster types emerged reliably at approximately 5 s post-stimulus onset and remained significant throughout the trial. These findings align with prior high-resolution fMRI evidence of robust motion direction selectivity in early visual areas (e.g., [19]).

**Fig 3.**
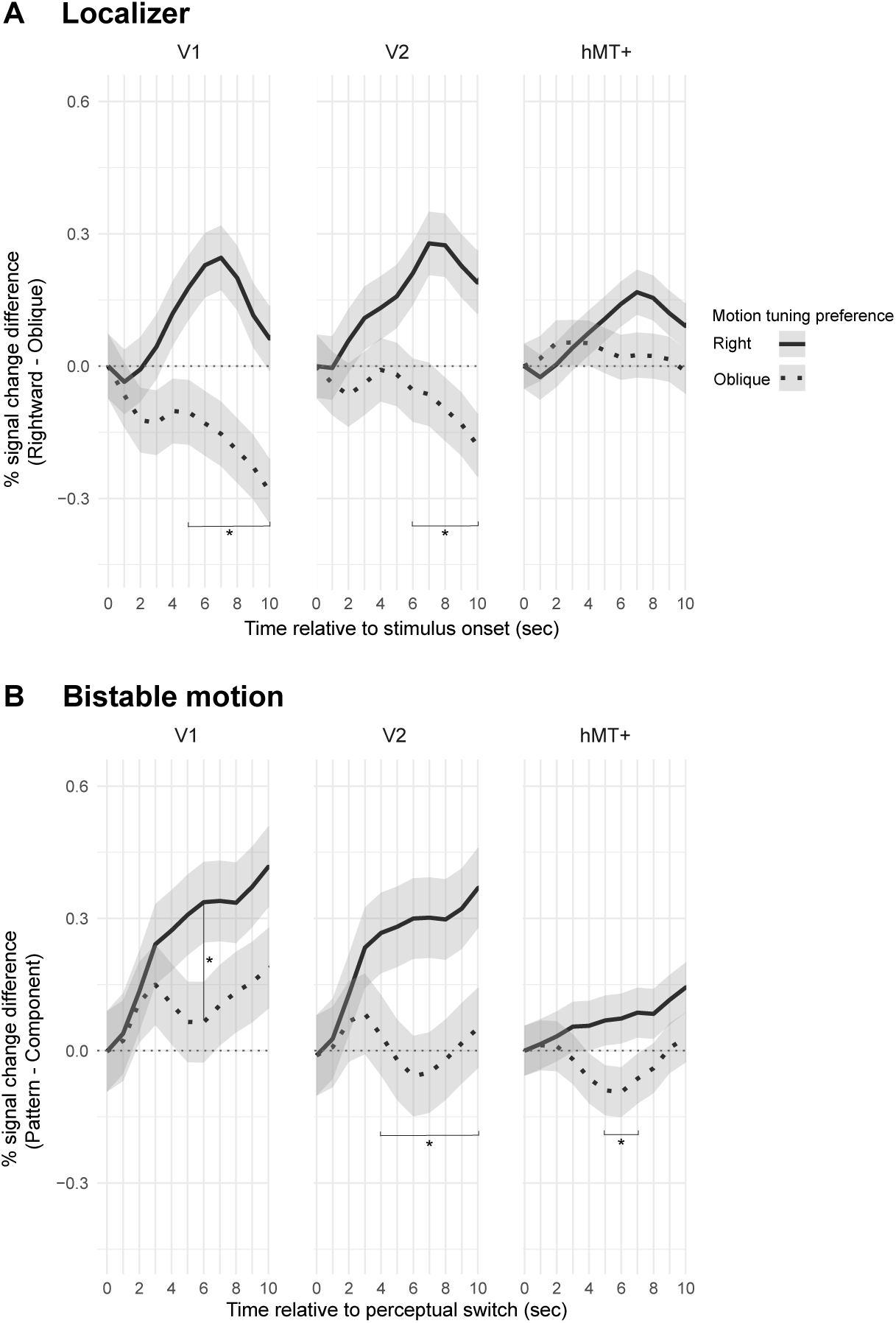
Event-related averages for both the localizer and the bistable motion stimuli. A. The temporal dependence of direction-selectivity (the difference in BOLD signals for rightward versus oblique motion) is shown for each ROI during the localizer motion sequence. Event-related average signal was time-locked to stimulus onset. Asterisks mark time points where post-hoc pairwise comparisons of estimated marginal means revealed a significant difference from baseline (Bonferroni-adjusted *p* < 0.05), confirming that direction-selective clusters preferentially respond to the physically presented motion direction. Shaded grey areas indicate standard errors. B. Pattern minus component signal change following a perceptual switch in the bistable stimulus is shown for each ROI. Event-related average signal was time-locked to perceptual-switch button presses. The increase in signal following a switch to the pattern percept was observed selectively in the cluster whose tuning matched the direction of the perceived motion.

In hMT+, rightward-selective clusters also tended to respond more strongly to rightward motion; however, ERA analyses did not reveal statistically significant direction specificity across clusters in this region. Importantly, direction selectivity in all areas was unequivocally confirmed using a GLM contrast across V1, V2, and hMT+, validating our functional definitions of the direction-selective ROIs (S1 Text, S2 Text and Table A in S3 Text).

For the bistable motion condition, the difference in ERA between pattern and component perceptual states is plotted over time for each direction-selective cluster in V1, V2 and hMT+ (Fig 3B). While direction-specific responses were generally weaker than in the localizer, ERA analyses revealed consistent divergence between rightward-and oblique-selective clusters. The difference in percent signal change between pattern and component percepts (pattern minus component) showed a reliable, selective increase in the rightward-selective clusters (solid lines) when subjects switched to the pattern percept. This event-related enhancement for the “preferred direction” (i.e., rightward) mirrors the selectivity observed in the localizer. The effect attained significance in V1, V2, and hMT+ with a peak ∼6 s after button presses, signaling percept onset.

In contrast, oblique-selective clusters (dashed lines) exhibited a more variable response profile following perceptual switches. ERA fluctuated between pattern and oblique preferential responses, despite these clusters being tuned more strongly for component motion directions. One possible explanation for this unstable direction-tuning responses in oblique-selective clusters is methodological: in a continuous bistable design, overlapping hemodynamic responses can produce temporal smearing that could obscure direction-specific effects in ERA analyses. Alternatively, the observed pattern may reflect genuine neural dynamics engaged during perceptual ambiguity.

### Eye movement directions do not differ between perceptual states

BOLD analysis demonstrated a differential brain response for pattern compared to component percepts, even though the driving physical stimulus remained constant. A potential alternative interpretation, however, is that the differential activity is not an effect of higher-level motion processing, but rather is driven by reflexive eye movements, known to covary with bistable visual stimulation [33]. In particular, if participants involuntarily moved their eyes along with the perceived motion direction, this would influence motion energy along that direction, introducing a bottom-up, stimulus-driven confound. To test this possibility, we recorded and quantified the direction of eye movements over the course of the experiment. We reasoned that if eye movements were entrained to the perceived motion direction, this should appear in the comparative distribution of eye movements around the fixation cross between the two states.

Even though subjects were instructed to fixate the central cross during the test period, the pattern of eye movement directions varied across observers (Fig 4A–C and S1 Fig) and did not necessarily align with the perceived directions of motion. Interestingly, the histograms of eye movement directions appeared to be very similar across perceptual states. Comparison of the distributions detected no statistical significance (LRT: *χ*^2^ = 31.1, *df* = 35; *p* = 0.66) (Table in S4 Text). Thus, this analysis excludes differences in oculomotor behavior as the basis of the percept-related BOLD effects.

**Fig 4.**
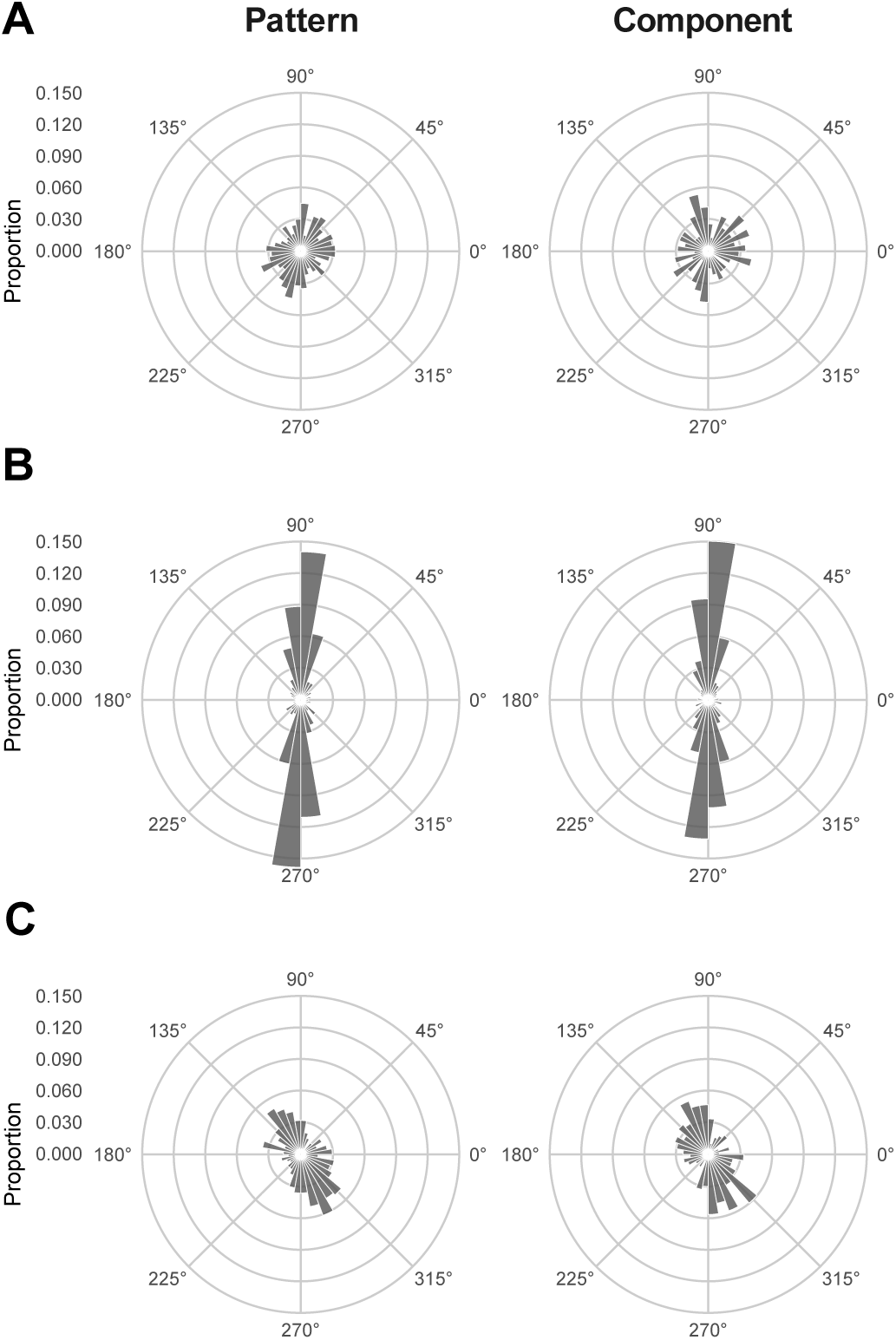
Distributions of eye movement directions for three observers during pattern and component states of perception. (A–C) Relative frequency distributions of eye movement directions for three observers, recorded during pattern (left) and component (right) states of perception. The scale to the right of A indicates the relative frequencies associated with each radius.

## Discussion

### Direction-selective voxels reflect perceptual decisions in visual cortex

We used a bistable motion paradigm to study whether changes in neural activity within three visual cortical areas were associated with perceptual state. Specifically, we asked whether perceptual state changes would be reflected in cortical areas as early as V1. In the bistable paradigm, the perception of two superimposed moving components spontaneously shifts between two transparent gratings sliding obliquely over each other (component state) or as a single plaid moving horizontally (pattern state), while the driving input stimulus remains unchanged. We found evidence that the activity in direction-tuned voxels was differentially modulated during component and pattern states as early as area V1 but also in areas V2 and hMT+. This demonstrates that perceptual representation is reflected across the extent of the visual cortical hierarchy that we investigated. Such perceptually driven activity in early visual cortex aligns with predictive coding frameworks in which higher-order areas send top-down predictions to lower-order areas, shaping sensory representations under ambiguity and biasing motion responses in early visual areas [16, 23], as supported by intracranial evidence that long term perceptual priors modulate feedback from higher order to early visual cortex during ambiguous perceptual states [34].

Specifically, direction-selective voxels in V1 tuned to the oblique directions showed no difference in response during the change in perceptual state while the voxels preferring the rightward direction displayed an increase in response during the pattern percept. These results support a circuit in which feedback input to V1 affects only the rightward direction-selective voxels. This pattern was reversed in areas V2 and hMT+ where there was a significant increase in response during the component epochs for oblique direction-selective voxels and no significant change in response for rightward preferring voxels. The peak difference in activity between the two states occurred approximately 5-6 s after the button press, consistent with the expected delay in hemodynamic response as previously reported with vascular space occupancy measures for ambiguous apparent motion, obtained using the motion quartet paradigm [19]. Importantly, the stronger perceptual modulation associated with the pattern state in V1, relative to V2 and hMT+, indicates a prominent contribution of top-down signals from higher-order areas to V1 in resolving visual ambiguity. The temporal dynamics of direction-selective responses further support this interpretation, suggesting a more complex mechanism in which rightward and oblique-selective clusters are differentially influenced by top-down signals. Under ambiguous motion, BOLD responses in areas V1 and V2 show reduced direction specificity and temporal profiles that converge with those in hMT+, consistent with enhanced top-down signals from higher-order areas to early visual cortex. This pattern aligns with findings from high-resolution fMRI showing that ambiguous moving stimuli elicit weaker direction-selective responses in V1 compared with unambiguous (i.e., physical) motion, and that the temporal dynamics between V1 and hMT+ become more similar during perceptual ambiguity [19]. Analysis of eye movements provided no evidence that they differentially affected the perceptual switching. Altogether, we conclude that the pattern-selectivity found in early visual areas reflects higher-level perceptual decisions rather than stimulus-driven differences.

Previous studies using animal models recording single neuron responses have demonstrated that at higher levels of the cortical hierarchy, there is an increase in the proportion of neurons showing pattern movement responses [4, 8–10], leading to the expectation of finding a stronger representation of the pattern percept in hMT+ than in areas V1 and V2. Paradoxically, we found the differential activation of the rightward cluster to pattern to be the highest in V1. Since visual input remained constant, the observed variation of activity across direction-selective units reflected internally-driven perceptual states and cannot be explained solely by the end-stopping property of neural units [35]. Although it is surprising that the pattern-selectivity was not significant higher in the hierarchy, especially in hMT+, it is possible that top-down processes are organized in chains, conveying signals towards lower-level areas, thereby resulting in an accumulation of signals at the lowest stage of the visual cortical hierarchy, area V1 [36, 37]. Such a proposition would be in accordance with the observation that top-down expectation effects are much stronger in area V1 than in areas V2 and V3 [38]. Importantly, the relative difference in voxel activity between the two different perceptual states, was significant in all the visual cortical areas investigated. This result is consistent with assumptions that pattern motion information is fed back to the earliest stages of cortical visual processing implying that visual motion perception reflects a global process operating across multiple levels of the cortical hierarchy, and that this cooperation—which is still poorly understood—requires interactions between both bottom-up and top-down processes [39].

### Role of top-down pathways in bistable motion disambiguation

While the current study does not allow unique identification of the higher-order source of the pattern motion signals in V1 and V2, convergent evidence suggests hMT+ as a likely candidate. For instance, recent ultra high field layer fMRI work shows that during ambiguous apparent motion, temporal coupling between hMT+ and V1 increases, and V1 responses reflect stronger top-down modulation compared with physical motion, supporting the notion that top-down signals contribute to percept stabilization [19]. Such laminar and temporal top-down signatures in V1 support the idea that early visual cortex activity is strongly influenced by top-down perceptual content rather than purely bottom-up input. Interestingly, results showing increased complex motion activity in area V1 are at odds with classical single unit recordings in macaque, which report that the vast majority of neurons respond primarily to the motion of the individual component gratings of a plaid stimulus, with only a sparse pattern-motion representation [4, 10]. One possible explanation has been that top-down signals primarily modulate population-level activity, rather than directly altering spiking output. Nonetheless, there is now evidence that top-down signals can also influence spiking activity in V1 in non-human primates [40, 41]. Another important consideration is that many single-unit recordings in monkey V1 have focused on neurons with foveal receptive fields, leaving open the possibility that top-down influences vary across the retinotopic extent of V1. Recent investigation of top-down signals to retinotopic subdivisions of areas V1 and V2 reveals important quantitative and also qualitative differences in the sources and strengths of projections to central and peripheral representations but also for the upper and lower hemifields [28].

In the present study we could not localize the complex motion response to specific retinotopic subdivisions of V1, and our experimental paradigm did not allow us to address how distinct patterns of top-down projections might impact these subdivisions. Future work combining high resolution fMRI with electrophysiological recordings will be needed to resolve discrepancies between hemodynamic and single-unit recording and to clarify the role of top-down processes across the full retinotopic map of V1. Top-down signals from hMT+ to V1 have been proposed to play a role in motion integration [15, 16, 42–45]. Moreover, Transcranial Magnetic Stimulation studies that disrupted hMT+ to V1 feedback transmission suggest a critical role of top-down signals in motion perception [46–49]. Further studies will be necessary to pinpoint the role of hMT+ on V1 in perceptual decisions associated with bistable motion perception. An important direction to pursue would be the investigation of the laminar profiles of activity using high intensity neuroimaging in V1, V2 and hMT+.

### Possible substrate for bistable motion

The perceptual switching with moving plaids differs from that in binocular rivalry. In binocular rivalry both input channels are stimulated (right and left eyes) and the percept fluctuates between that evoked by each of the input channels (eyes) [50–52]. With the moving plaids, as presented here, the input is a pair of obliquely oriented gratings that presumably stimulate cortical units tuned selectively to the direction of motion of these two components (oblique). The percept, however, fluctuates between one corresponding to the tuning of the oblique components and a rightward pattern motion whose direction is presumably outside the bandwidth of the oblique-selective units. This characteristic of the design supports the hypothesis that a circuit for the pattern motion requires a hierarchical model in which the responses of univariant, narrow-bandwidth sensors at an early processing stage, such as V1, are combined to extract the pattern motion at a second processing stage, such as MT, evidence for which was reported in the initial physiological experiments using moving plaid stimuli [53].

Is the pattern of differential responses observed across direction-selective clusters and cortical areas consistent with known physiological properties of neuronal responses to moving plaid patterns? In macaque area V1, neurons identified through antidromic stimulation as projecting to pattern-selective cells in MT have been characterized as direction-selective complex cells mostly in layer 4B that respond selectively to component motion rather than global pattern motion [53]. Although fMRI voxel responses reflect population-level activity, they are likely biased toward the neuronal clusters preferentially activated by the localizer. In our case, the oblique motion localizer—composed of oblique dashed-line segments moving unambiguously in two oblique directions—is expected to preferentially recruit component-selective complex cells as described above for V1. The absence of a significant differential response between perceptual states in V1 oblique-selective voxel clusters is therefore consistent with the hypothesis that these neurons are not influenced by percept-related top-down signals. In contrast, the rightward motion localizer, also composed of oblique dashed-line segments was unambiguously perceived as moving rightward. This stimulus is unlikely to activate rightward-selective classical component cells in V1 since it contains no vertically oriented components. Instead, we propose that this stimulus either preferentially engages V1 end-stopped neurons sensitive to rightward motion direction defined by line terminations located in layers 2/3 [35, 54–57] and/or non-orientated direction-selective neurons known to be located in layers 4 and 2/3 [58–61]. To explain the strong differential responses observed in rightward-selective V1 voxels across perceptual states during bistable motion, we propose that these signals reflect top-down modulation of end-stopped and/or non-oriented neurons linked to perceptual interpretation. This prediction could be tested using high-field laminar-resolution fMRI.

In macaque MT, direction-selective neurons are characterized as either component or pattern direction-selective [4, 62, 63]. Interestingly, in human hMT+, we found that in spite of a significant differential effect across perceptual states between oblique- and rightward-preferring voxel clusters, the pattern shifted from that shown in V1. Oblique-selective voxels were significantly activated during component perceptual states while no significant differential activation occurred in rightward-selective voxels across the two perceptual states. Assuming that the oblique localizer preferentially recruits component direction-selective populations, the increased activation during component perceptual states suggests that their activity is relatively suppressed during pattern states. Conversely, the absence of modulation in rightward-selective voxels implies that the underlying neuronal populations are not differentially engaged across perceptual states (also, see [63]).

This interpretation is consistent with computational models of pattern direction-selective neurons in macaque MT, which propose that their selectivity arises from the integration of a broad range of orientation inputs, shaped by a balance of excitation and inhibition across orientation channels [62]. In these models, pattern selectivity is not explained by a simple elevation in baseline activity of pattern-selective neurons, but instead arises from the suppression of responses tied to individual component motions (motion opponency) together with broad pooling and direction-tuned normalization that preferentially supports coherent pattern motion over isolated component directions. Area V2 exhibits an intermediate response profile, partially resembling hMT+, suggesting that top-down modulation occurs in a hierarchical chain, in which feedback from higher-order areas shapes activity progressively across the visual cortical hierarchy. Thus, one can envision a circuit in which hMT+ pattern-selectivity arises from the integration of bottom-up input from V1 neurons, including end-stopped cells responding to plaid intersections, while pattern-selectivity in V1 reflects top-down modulation from hMT+, enhancing activity in end-stopped direction-selective populations. Periodic adaptation may tip the balance between perceptual states, with a relatively slow time constant controlling the build-up of adaptation [43, 64].

## Conclusion

We find that perceptual switches in motion perception are accompanied by specific brain states that are present across multiple levels of the visual cortical hierarchy. Direction-tuned voxels in V1, V2, and hMT+ reliably differentiated pattern and component percepts; notably, the magnitude of pattern-selective modulation was greatest in V1. This finding aligns with the importance of top-down signals in relaying high-level contextual information to low-level visual areas in order to interpret visual input [65] or to generate internally-driven mental images [29, 30]. Furthermore, our results provide evidence that perceptual states were associated with spatially distinct direction-selective clusters as early as area V1. This suggests that distinct neural units encode concurrent perceptual states and that the spatial segregation of information is preserved across hierarchical levels. The fact that we and others observe abundant pattern-selectivity in area V1 underscores the critical role of top-down connectivity in motion perception and indicates that perceptual experience emerges from dynamic interactions across multiple levels of the visual cortical hierarchy.

## Materials and methods

### Ethics statement

The experimental procedure was covered by the CMO regio A-N 2014/288 blanket approval, “Imaging Human Cognition”, between the Donders Center for Cognitive Neuroimaging (DCCN) and the Medical Research Ethics Committee.

### Subjects

Thirty-seven healthy subjects (25 females, mean age 25.5, sd=3.71) with normal or corrected-to-normal vision were recruited for this fMRI study. All participants provided informed consent and received 20 euros for their participation.

### Experimental design

#### pre-fMRI training

Before entering the scanner, each participant practiced the task during a short behavioral training period in order to ensure that they experienced the spontaneous perceptual switching during the bistable motion task. This also provided practice in reporting the perceptual shifts and fixation training.

#### Localizing motion- and direction-selective regions

In the same session as the main experiment, 3 runs using the functional localizer were performed. The localizer served two main goals: first, defining bilateral motion-selective regions in V1, V2 and hMT+, and secondly, selecting within each region rightward- and oblique-selective clusters that matched the perceived directions of the two perceptual states during the bistable motion task.

The functional localizer consisted of short trials (20 seconds) of single gratings moving unambiguously as well as static ones (control condition) interleaved with ten-second inter-trial intervals. The stimulus motion was designed specifically to be in the direction matching either the perceived component directions (oblique - upward and downward) or the pattern direction (rightward direction) (Figure A in S1 Text and S1 Video). Each condition was repeated 13 times per run. In order to induce an unambiguous motion percept while keeping unchanged the design of the oriented lines moving behind the aperture, we used dashed-line gratings, allowing the extraction of information from the local edges of the dashes and thereby restoring a two-dimensional motion signal. We also enhanced the contrast of the stimulus (white lines moving on top of a black background), thereby ensuring a strong and unambiguous perception of motion direction.

#### Inducing a bistable motion perception

Observers fixated a central cross while viewing a bistable motion stimulus consisting of two non-collinear, overlapping square-wave gratings (plaids). Each grating had a spatial frequency of 0.32 cycles/deg and an unequal duty cycle square-wave texture 77.8% high contrast phase and 22.2% low contrast phase). Component gratings moved orthogonally to their orientation at a speed of 0.63 deg/s, with orientations set at ±65 deg relative to horizontal (130 deg separation). Both gratings were defined with identical contrast values (0.7 in PsychoPy’s normalized RGB contrast space). Transparency was induced by assigning one component grating reduced opacity (0.2). These parameters were chosen based on prior studies of plaid motion bistability [66–68] and preliminary psychophysical testing (N = 5 observers) to optimize perceptual alternations. The stimulus was presented within a circular annular field (outer border: 10 deg, inner: 1.3 deg), with smoothed edges (raised cosine for outer, Gaussian for inner) and a central fixation cross subtending 0.5 deg. The background luminance matched the space-averaged luminance of the gratings (CIE *xyY* = (0.33, 0.34, 100 cd/m^2^)).

To bias perception toward the pattern or component state, the contrast of plaid intersections was manipulated [67, 69–72]. Plaids with enhanced intersections tend to induce the pattern percept, whereas reduced intersection contrast favors the component percept. In the present study, we used a hybrid plaid design in which one grating had reduced opacity, limiting perceptual states to two: a component state with a fixed depth-ordering (darker grating on top) and a pattern state moving rightwards (Fig 1A and S2 Video).

Extended viewing of the plaid stimulus induced bistable perception, with participants alternating between the two mutually exclusive states. Participants reported perceptual switches dynamically via a bimanual 8-button response pad (HHSC-2×4-C), indicating components, pattern, or mixed percepts. Component and pattern responses were assigned to separate hands to minimize confusion and optimize response times. Subjective reports were later used to define experimental conditions and align fMRI and eye-tracking data. To ensure reliable measurement of event-related BOLD responses, percepts shorter than 2 s or longer than 30 s, as well as mixed reports, were excluded (Zhang et al., 2017). On average, these exclusions accounted for 11.3% of stimulation time (95% CI = [7.4, 15.2]). Observers completed five runs, each consisting of six 1-minute trials separated by 10-second inter-trial intervals. The transition rate between perceptual states served as a measure of motion bistability, and roughly equal durations for component and pattern states were expected. Participants failing to reliably experience bistable perception were excluded (two subjects).

### Imaging data acquisition

All MRI data were acquired on the 3T MAGNETOM Skyra MR scanner (Siemens AG, Healthcare Sector, E rlangen, Germany) at the Donders Center for Cognitive Neuroimaging (Nijmegen, Netherlands), using a product 32-channel head coil. All participants performed a single fMRI scanning session of approximatively 1 hour. The structural MRI sequence, a T1-weighted Magnetization Prepared Rapid Acquisition Gradient Echo (3D-MPRAGE, 256 sagittal slices, TR = 2300 ms, TE = 3 ms, flip angle = 8 deg, resolution: 1mm ISO), was acquired together with a head scout and localizer (32-channel head coil). Furthermore, an initial slice positioning fieldmap was collected to ensure the functional coverage of V1, V2 and hMT+ (66 coronal slices). The functional MRI acquisition consisted of Multi-Band accelerated Echo Planar Imaging (EPI) Pulse sequences (MB6, 66 coronal slices, TR = 1000 ms, TE = 34 ms, Anterior-Posterior phase encoding, resolution: 2mm ISO). This type of accelerated sequence has the advantage of using a short repetition time and a 6-fold acceleration factor (number of slices simultaneously excited) that results in the acquisition of a multiband signal in a single EPI echo train. The use of multiband-accelerated MRI sequences allowed shorter repetition times than conventional sequences, increasing temporal sampling density and enabling more accurate capture of rapid neural changes associated with perceptual alternations [73]. A sequence with the inverted phase encoding gradient (Posterior-Anterior) was conducted to efficiently correct for EPI gradient field non-linearity distortions. Imaging data analyses were implemented via Nipype v1.8.2 interface (combining SPM, FSL and Freesurfer modules).

### Preprocessing steps

Preprocessing steps were applied following the fMRIprep analysis pipeline. First, T1w images were corrected for intensity non-uniformity [74, 75] and skull-stripped (Nipype implementation of the ANTS atlas-based brain extraction workflow). Brain tissue segmentation of CerebroSpinal Fluid (CSF), White-/ Grey-Matter (WM/GM) was defined (based on FSL Fast toolkits). Volume-based spatial normalization of the T1w reference was performed to the MNI152NLin2009cAsym standard space through nonlinear registration (ANTS registration tool; [76]).

Regarding BOLD series, a reference volume was computed out of the aligned single-band reference images (SBRefs). A field map estimating susceptibility distortions was estimated based on the reversed phase encoding EPI sequence (AFNI; [77]) and enabled unwarping the data, providing a corrected EPI reference. This latter was used to co-register functional data to the T1w reference (using bbregister in Freesurfer) by implementing a boundary-based registration [78] with 6 degrees of freedom. Head-motion parameters were estimated with respect to the functional reference (transformation matrix including 6 rotation and translation parameters, using mcflirt in FSL described in [79] before any spatiotemporal filtering was applied. The BOLD signal was slice-time corrected (using 3dT-shift in AFNI; Cox1996). The resulting BOLD time-series were then resampled to both native and standard space by applying a single composite transform to correct for head-motion and susceptibility distortions.

FMRIPrep calculates several confounding time-series based on the preprocessed BOLD signal. Framewise displacement (FD) and the derivative of the root mean square variance over voxels (DVARS) were calculated for each functional run, both using their implementations in Nipype (following the definitions by Power et al., 2014). FD reflects the instantaneous head motion between consecutive volumes, based on the summed changes in head position parameters, while DVARS quantifies the rate of change in BOLD signal intensity across the brain between successive frames.Three global signals were extracted within the CSF, the WM, and the whole-brain masks. Additionally, a set of physiological regressors were extracted to allow for component-based noise correction using the CompCor method [80]. Principal components were estimated after high-pass filtering the preprocessed BOLD time-series (using a discrete cosine filter with 128 s cut-off) for the two CompCor variants: temporal (tCompCor) and anatomical (aCompCor). tCompCor components are calculated from the top 2% variable voxels within a mask covering the subcortical regions. aCompCor components are calculated within the intersection of the aforementioned mask and the union of CSF and WM masks calculated in T1w space, after projection to the native space of each functional run.

After preprocessing with fMRIPrep, the confounds were inspected to determine if data met the criteria for inclusion. Functional runs were excluded if more than 30% of time points exceeded a FD of 0.5mm. This led to the full exclusion of one subject and of two runs of bistable motion for another subject. For the remaining subjects, we included confounding factors in the estimation of BOLD time-series, namely the six head-motion parameters, the first six aCompCor components from WM and CSF mask, and FD.

BOLD data were smoothed using SUSAN (Smallest Univalue Segment Assimilating Nucleus) nonlinear noise reduction filtering [81], which reduces noise while preserving underlying structure by averaging each voxel only with neighboring voxels of similar intensity (i.e., within the same local region) rather than uniformly across space. This technique preserves edges and anatomical details better than linear filters by limiting smoothing to voxels whose intensities lie within a defined brightness threshold. A 6 mm FWHM Gaussian equivalent kernel was applied, and the SUSAN brightness threshold was set to 75% of the median image intensity for each run.

Subsequently, each volume of every run was scaled so that the median value of a specific run was set to 10000, and a scaling factor was estimated for intensity normalization. Finally, BOLD data were band-pass filtered using high-pass (0.008 Hz) and low-pass filtering (3^rd^-order polynomial function using the method of linear least squares) (see [82, 83]).

### Regions of interest

#### Regions of interest (1): Motion-selective regions within visual cortex

The Freesurfer reconstruction tool enabled delineating anatomical regions of interest bilaterally, namely retinotopic visual areas V1 and V2, as well as hMT+ complex, the contiguous region of gray matter in the posterior middle temporal region responding more strongly to moving gratings than to stationary ones (selection of the 100 most specific voxels from the [motion — static] GLM contrast, *p* < 0.01). For each participant, we first extracted bilateral V1, V2 and hMT+ masks (average size±sd, isotropic resolution 1 mm: 9946.6 ± 753.1 voxels; 17037.0 ± 953.9 voxels; 4060.5 ± 388.2 voxels, respectively). Within these anatomical masks, we used a functional localizer to reveal direction-selective populations of voxels (see example in FIgure B in S1 Text).

#### Region of interest (2): Localization of direction-selective regions

Each area’s average response to the unambiguous localizer sequence was used to map direction-selective clusters independently of the bistable perception experiment. Prior to determining the optimal cluster size, we considered how generalizable direction-tuning was across localizer runs. To evaluate this, we used a 3-fold cross-validation approach where each iteration consisted of two steps: 2 runs were picked to select the N most direction-selective voxels, and the remaining run was used to test the specificity of the selection in the estimated GLM contrast [rightward - oblique]. This leave-one-run-out approach was repeated for different cluster sizes (N = [25, 50, 75, 100] voxels) (**??**). This range of sizes was determined based on the smallest functionally defined region hMT+. Quantitative analyses of optimization of unilateral hMT+ functional localization reported an average size that is equivalent to 100 to 150 voxels (for a voxel size of 2 mm^3^) [84]. Therefore, the maximal cluster size for both rightward and oblique conditions was set to be less than or equal to half of the hMT+ minimal surface (i.e., 50 voxels per hemisphere, or 100 in total), and superior to one quarter of the total surface to ensure sufficient signal-to-noise ratio (*>*25 voxels). Six participants were excluded from the group level analysis because no significant direction-selective clusters emerged from the localizer. In two additional participants, only the hMT+ data were discarded — the remainder of their data being retained — because the direction-selective hMT+ clusters were either too small and/or failed to reach the GLM contrast threshold.

### Statistical Analyses

#### Neural correlates of bistable motion perception

BOLD responses were assessed by applying an ROI-based fixed-effect general linear model (GLM) approach using the FSL implementation in Nipype. All regressors representing experimental conditions were convolved with a Double-Gamma hemodynamic response function (dgamma-HRF). The first-level (run-level) design matrix included predictors (experimental condition, button press) and motion correction parameters as confound predictors. Voxel timeseries were z-transformed and corrections for serial correlation were applied, using a first-order autoregressive estimator. The second-level (or subject-level) GLM design matrix was then created for each subject by merging the run-based contrast estimates.

For the localizer sequence (3 runs), three predictors (oblique and rightward motions, static stimulus) were represented with boxcar functions. The average BOLD activity (*z*-score) was then contrasted to localize [rightward - oblique] direction-selective clusters within the early visual cortex.

For the bistable motion sequence (5 runs), we modeled the perceptual report (component, pattern and mixed percepts) with Single Impulse (or stick) functions. Within the previously defined direction-selective clusters, we mapped the main contrast of interest, pattern minus component.

For each of the 28 included observers, we estimated the difference in BOLD activity between the two perceptual states, pattern and component within rightward and oblique clusters (Table B in S3 Text). Our main hypothesis was that the pattern state would preferentially elicit response in rightward clusters, whereas the component state would more strongly activate oblique clusters. In order to test this hypothesis (against the null hypothesis of no preferential activation of clusters with percept), we tested the difference of pattern versus component BOLD contrast between direction-tuned clusters, i.e., the interaction between the preferred perceptual state and the region’s direction-selectivity.

#### Linear Mixed Effect model

Group-level statistics were calculated by implementing linear mixed-effects models (LME) with the **lme4** package [85] in the R statistical environment [86] in RStudio (v2022.07.1+554) to account for between-subject variability, while testing the effect of variables (or interactions of variables) that best fitted the measured response variables as described in S2 Text. This model enabled us to test the main hypothesis that there is a significant interaction between the direction-selective BOLD activity and the neural correlates of perceptual decision in early visual cortex. Statistical significance of fixed effects was assessed using nested likelihood ratio rests. This procedure provides an appropriate estimation of inferential statistics for mixed-effects models with hierarchically nested models. Reported z-values were obtained post-hoc using the glht function in the **multcomp** package [87] and include corrections for multiple tests. Note that z-values obtained from post hoc comparisons differ from those derived from individual GLM contrasts.

#### Event-related Averages

Within direction-selective clusters defined from preprocessed contrast maps (preferring oblique or rightward motion), we extracted BOLD signal time courses to examine the temporal dynamics of physical motion (from the localizer data) and perceived motion (from the bistable stimulus) across the visual cortical hierarchy. For the localizer, BOLD signal was time-locked to trial onset and averaged separately for each motion direction (rightward and oblique) within each direction-selective cluster. For the bistable motion dataset, event-related responses were time-locked to button presses, and averaged separately for the pattern and component conditions within direction-selective clusters. Events with durations ≤ 10 seconds were excluded to ensure comparability between the localizer and bistable motion datasets. Voxel-wise time-series were temporally smoothed using a Savitzky–Golay filter (polynomial order 3, frame size 5) to reduce noise while preserving hemodynamic features.

For each event, BOLD signals from all voxels within each ROI mask were extracted over a temporal window beginning at trial or percept onset (*t*_0_) and extending for 10 TRs (11 volumes). Baseline activity was defined as the voxel signal at *t*_0_, and percent signal change was computed at each subsequent time point as (*t* − *t*_0_)*/t*_0_ × 100.

Because the perceptual bistability paradigm elicits closely spaced perceptual events without inter-stimulus intervals, we focused on relative differences between direction-selective clusters over time rather than absolute percent signal change values. To assess direction selectivity in the localizer, activity during rightward and oblique motion stimuli was contrasted (rightward – oblique). To examine how perceptual state modulated direction-selective responses in the bistable dataset, activity during pattern and component percepts was compared (pattern minus component). For each ROI and time point, differences in direction-specific BOLD responses between direction-selective clusters were evaluated using estimated marginal means, followed by Bonferroni-corrected pairwise comparisons across time points to control for multiple comparisons.

#### Analysis of microsaccadic eye movements

An Eyelink 1000+ setup synchronized to the fMRI scanner tracked the oculomotor activity (raw eye position, saccades, blinks, pupil dilation) throughout the session. The Eyetracker system was set on monocular mode (tracking left eye), and a 9-dot calibration was performed before data collection began, to ensure high gaze accuracy and quality of recording. Eyetracking constituted an important measure to obtain during the whole session, as it has been shown that in the absence of an attentional/fixation task, eye movements reliably predict perceptual state [88]. Moreover, although observers were trained to fixate on a central target, involuntary eye movements persisted in response to a moving stimulus. Therefore, we analyzed the recordings of the individuals included in the fMRI group-level analyses while discarding the data of five participants for whom the eyetracker encountered technical problems during the session, hence resulting in an eye-tracking dataset of 23 observers (S1 Fig).

Raw gaze data (x / y screen coordinates) were first converted from screen pixel units into visual angle in degrees. We then filtered the data to exclude large saccadic eye movements, in order to focus on microsaccades [88]. Following the guidelines of the EyeLink system, we applied exclusion criteria similar to the saccadic thresholds defined in the EyeLink1000 User Manual (velocity > 40 deg/s or acceleration >8000deg/s^2^) [89]. In addition, we discarded discontinuous segments shorter than 50 ms to avoid artefacts. For each subject and each run, we computed the mean gaze position (horizontal and vertical) during fixation periods — when participants were instructed to maintain steady fixation — to serve as a run specific baseline (“center”). Gaze positions were then binned on a 32 × 32 2D spatial grid spanning −5 deg to +5 deg in both horizontal and vertical dimensions around the fixation center. Each valid sample was assigned to the appropriate bin, and the number of samples per bin was counted separately for each perceptual condition. Finally, for each condition, bin counts were normalized by the total number of valid gaze samples to yield a relative frequency distribution of gaze positions. We then derived a “movement vector” between successive segments by computing the difference in mean positions (Δ*x,* Δ*y*) from one segment to the next, and converted this into a movement direction (angle, in degrees) and movement amplitude (Euclidean distance). We binned the resulting movement directions into 10 deg angular bins, and for each perceptual condition we calculated the relative frequency of movement directions (normalized counts per bin) to obtain a directional distribution of gaze shifts.

Histograms of eye movement direction counts, across perceptual states, were compared with a Generalized Linear Mixed-effects model (GLMM) with a Poisson family (default log link), implemented with the glmmTMB function from the **glmmTMB** package [90] in R. A subject-level random effect for eye movement direction was included to account for differences in eye movement distributions between subjects while testing within-subject differences between perceptual states. Model fits including and excluding a fixed-effect interaction between perceptual state and eye movement direction were compared using a nested likelihood ratio test. To assess the possibility of overdispersion, we calculated the ratio of the sum of squared Pearson residuals from the model fit to the residual degrees of freedom (number of observations minus the number of estimated parameters). Under the Poisson assumption, this ratio is expected to be approximately 1, with substantially larger values (> 2) indicative of overdispersion. The value obtained was 0.63, not supporting the hypothesis of overdispersion

## Supporting information

**S1 Text. Mapping direction-selective subdomains.**

**S2 Text. Localization of direction-selective subdomains.**

**S1 Fig. Eye-movement directional profiles during pattern and component perceptual states.** (A-T) Radial plots of eye movement direction frequencies for participants 3, 4, 5, 11, 12, 13, 14, 15, 16, 18, 19, 21, 27, 28, 29, 31, 32, 34, 35 and 36, illustrating the distribution of eye movement directions.

**S3 Text. Tracking perceptual decision within direction-selective clusters.**

**S4 Text. Testing differences in eye movement direction histograms between perceptual states.**

**S1 Video. Short examples of the video sequences used to localize oblique-and right-preferring voxels**

**S2 Video. A video sequence representing the plaid stimulus composed of 2 obliquely drifting gratings.** When steadily fixating the central cross, most observers experience spontaneous shifts between perceiving two transparent gratings sliding across each other and a single coherent plaid moving rightward. Because this video does not reproduce the calibrated display properties and controlled viewing conditions of the experiment, it may not faithfully replicate the perceptual effects reported in the article.

## Acknowledgments

We thank Paul Gaalman for his expert help with scanning and the members of the Predictive Brain Lab for their input during the development of this project.

## Abbreviations

BOLD: Blood Oxygen Level Dependent
CI: Confidence Interval
CSF: Cerebrospinal Fluid
DVARS: derivative of the root mean square variance over voxels
EPI: Echo planar imaging
ERA: Event-related Average
FD: Framewise displacement
fMRI: functional Magnetic Resonance Imaging
GLM: General Linear Model
GLMM: Generalized Linear Mixed-effects Model
GM: grey matter
hMT+: human medial temporal complex
LRT: likelihood ratio test
LME: Linear Mixed-effects model
MT: medial temporal area
MST: medial superior temporal area
ROI: Region of Interest
SUSAN: Smallest Univalue Segment Assimilating Nucleus
TE: Time to Echo
TR: Repetition Time
WM: White Matter

