## Supplemental Text 1 for "Top-down perceptual modulation of pattern but not component tuned motion circuits in V1"

#### **Supplementary Text S1:** Mapping direction-selective subdomains

Within each visual area (V1, V2 and hMT+), we identified the top 25 voxels tuned to rightward and oblique motion based on the average contrast estimate from the subject‑level GLM across all localizer runs ([rightward – oblique]). Subdomains exhibiting strong direction tuning (positive *z*‑scores for the rightward subdomain; negative *z*‑scores for the oblique subdomain) were then defined as regions of interest for the group‑level analyses in the bistable motion experiment. Supplementary Figure S1 illustrates an example of these direction‑selective subdomains mapped onto the cortical midthickness surface for a representative participant.

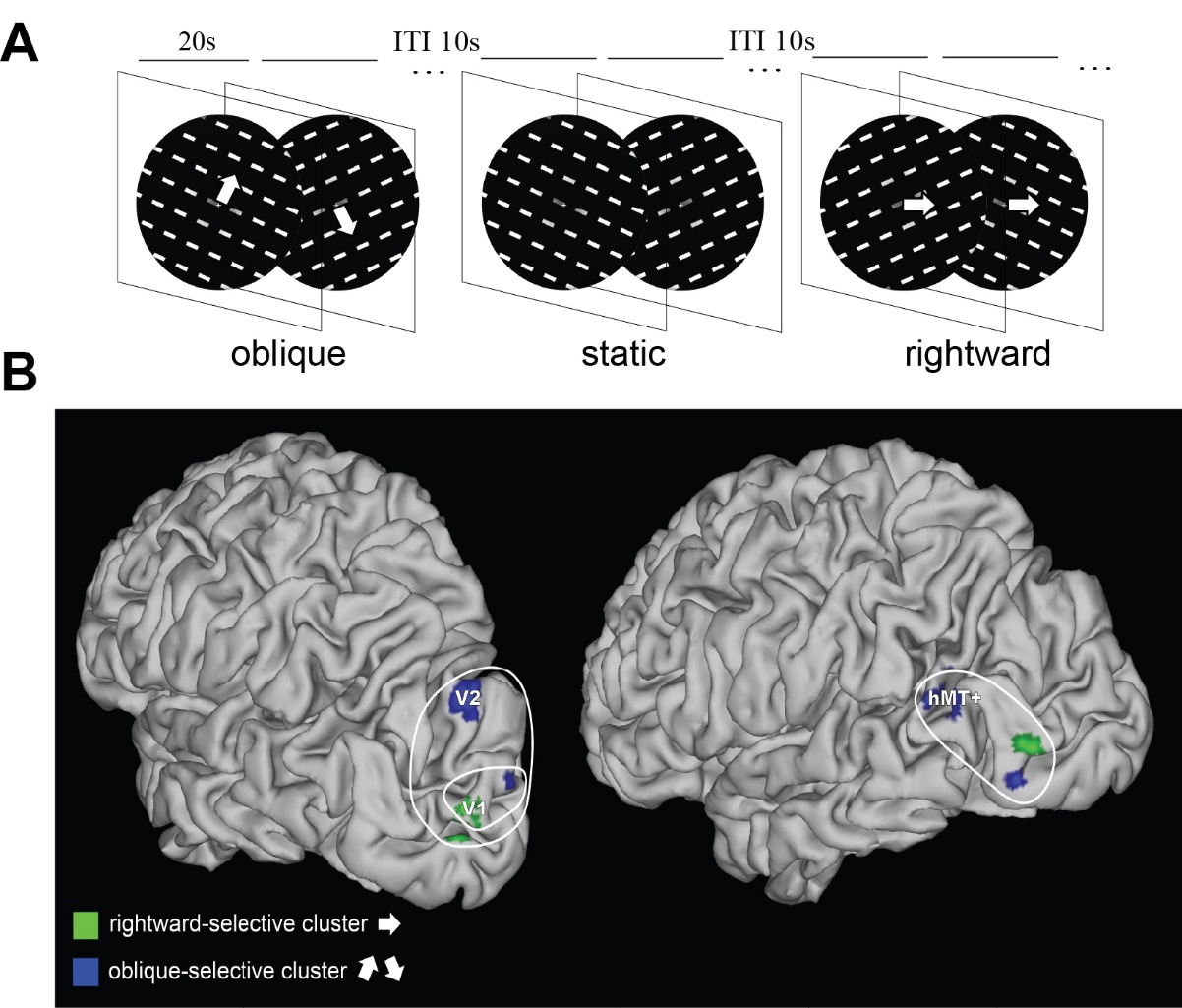

*S1 Figure: Mapping direction-selective subdomains in visual areas V1, V2 and hMT+ complex*

**A.** *Dashed-line gratings moving either rightward or obliquely were shown to participants during 20-second trials. Gratings orientation and direction matched those perceived in the bistable plaid stimulus. Based on this independent localizer, direction-selective GLM contrast estimates enabled to define direction-selective units within visual cortical areas.*

**B.** *Left view represent rightward (in green) and oblique (in blue) top direction-selective voxels within areas V1 and V2. Right view shows the corresponding direction-selective clusters within hMT+. Example: Subject 21, left hemisphere*
