## Supplemental Text 2 for "Top-down perceptual modulation of pattern but not component tuned motion circuits in V1"

#### **Supplementary Text S2:** Localization of direction-selective subdomains

In order to assess the generalization of direction-selectivity across localizer runs, a 3-fold cross-validation test was performed to select and estimate contrast [rightward - oblique] on distinct runs. The average z-score was then calculated for different sizes of direction-selective subdomains (N = [25, 50, 75, 100] voxels). The effect of the subdomain size was evaluated based on a linear mixed-effects model (LME) (Bates et al., 2015) using the *lmer* function in the **lme4** package of R with the following expression for the formula object:

$$z \sim Area/Size/SubDom + (1 | Sub) + 0,$$

where $z$ represents the average z-score of the [rightward – oblique] GLM contrast estimate conditional on the subject random intercept term, (1 | Sub) assumed to be normally distributed with mean 0 and estimated variance, $Area/Size/Subdom$ is the nested fixed effect of Subdomain (Subdom: rightward- or oblique-selective) within size treated as a factor level (Size: 25, 50, 75, 100) within each visual area (Area: V1, V2, hMT+). The 0 added to the end excludes the intercept and has the effect that the 3-way interactions correspond to the differences between subdomain for each size and within each area, which are the quantities of

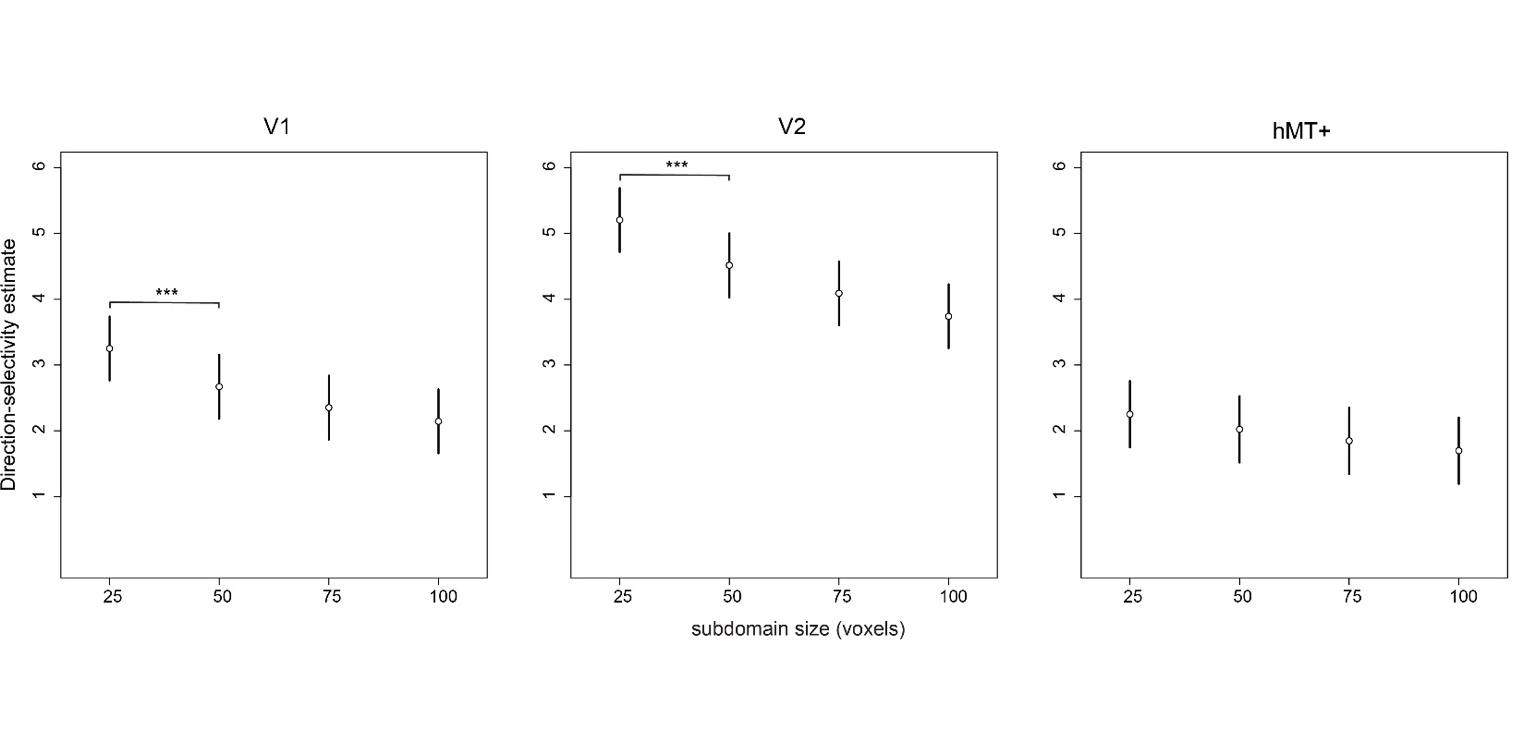

S2 Figure 1: Direction-selectivity of subdomains is consistent across individual runs

Mixed Effect model estimates of direction selectivity (normalized contrast of BOLD activity: [rightward – oblique]) within motion-specific regions of early visual areas as a function of subdomain size (from 25 to 100 voxels). The error bars indicate 95% confidence intervals. The selected subdomains were specific to direction, and the effect generalized across runs (Z>0, **p< .01**). In V1 and V2, the smallest size (25 voxels) maximized direction selectivity showing a significantly higher selectivity (V1: Z= 3.25; V2: Z=5.17, *** **p< .001**). In hMT+ subdomains, although direction selectivity were less consistent across runs, and not as clearly dependent on the size. However, the responses attained statistical significance (hMT+ (25 voxels): Z=2.25, **p< .01**)

N= 28 observers.

interest to test.

The fixed-effect estimates from the model fit are plotted as a function of subdomain size. Overall, the direction-selectivity was maximized for N = 25 voxels (the smallest size) (the null hypothesis of no difference in selectivity explained by subdomain size was rejected at p < 0.001) (Supplementary Figure S2).
