## Supplemental Text 3 for "Top-down perceptual modulation of pattern but not component tuned motion circuits in V1"

#### **Supplementary Text S3:** Tracking perceptual decision within direction-selective clusters

Average direction-selective responses per subdomain:

**A** **Localizer**: [Rightward – Oblique] contrast estimate shows high direction-selectivity regarding rightward (z-score ± 95% CI > 0, **p< .001**) and oblique (z-score ± 95% CI < 0, **p< .001**) subdomains across visual areas at the group-level.

**B** **Bistable motion**: [Pattern – Component] contrast estimate shows high specificity regarding rightward (z-score ± 95% CI > 0, **p< .001**) and oblique (z-score ± 95% CI < 0, **p< .001**) subdomains across visual areas at the group-level.

N=28 observers

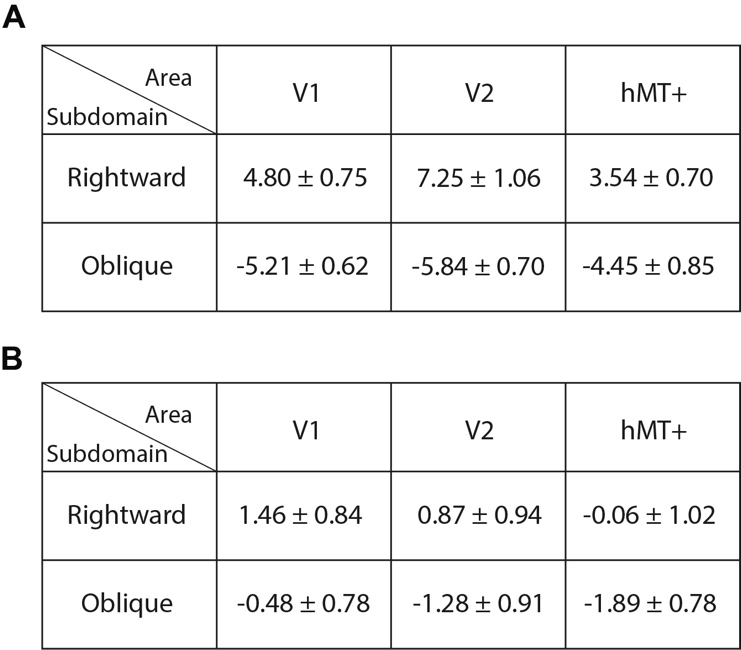

The tables above summarize the group average z-scores (± 95% confidence intervals) for the localizer contrast [rightward – oblique] (Supplementary Table S1.A) and the bistable motion contrast [pattern - component] (Supplementary Table S1.B). The contrasts are estimated per area (columns: V1, V2, hMT+) per subdomain (rows: rightward and oblique).

If the previously localized direction-selective subdomains reflect perceived motion, we expect the same trend in the z-scores representing pattern/component selectivity than that of the localizer (Z > 0 in rightward subdomain and Z < 0 in oblique domain). Our results are consistent with this prediction.
