## Supplemental Text 4 for "Top-down perceptual modulation of pattern but not component tuned motion circuits in V1"

#### **Supplementary Text S4:** Testing differences in eye movement direction histograms between perceptual states.

The eye movement histograms between perceptual states were compared using a Generalized Linear Mixed-effects model with a Poisson family and the default log link, using the function glmmTMB from the glmmTMB package in R. Two models were fit: one that included an effect of perceptual state and one that did not. The formula objects for these two models are

M1: Counts ~ Angle * Percept + offset(log(TotalCounts)) + (1 | Subject:Angle)

and

M0: Counts ~ Angle + Percept + offset(log(TotalCounts)) + (1 | Subject:Angle),

where Counts is the number of eye movements that fell into each direction (Angle), Percept is 2-level factor indicating the perceptual state and TotalCounts the total number of eye movement directions measured in a given state for each subject. Since the default log link is used, the offset term normalizes the counts by the total number of eye movements counted for a given subject in a given direction. The random intercepts for each combination of Subject and Angle are assumed to be independently and identically distributed and drawn from a normal distribution with mean 0 and variance estimated in the fitting procedure. The two models are compared with a nested likelihood ratio test. The results are shown in the table below.

Nested Poisson GLMM comparison testing the interaction between eye movement direction and perceptual state :

|  | df | AIC | BIC | logLik | deviance | $\chi^{2}$ | df | p |
| --- | --- | --- | --- | --- | --- | --- | --- | --- |
| M0 | 38 | 8522.1 | 8726.9 | -4223.1 | 8446.1 |  |  |  |
| M1 | 73 | 8557.5 | 8950.9 | -4205.8 | 8411.5 | 34.636 | 35 | 0.4856 |
