## Supplementary figures and images for "Top-down perceptual modulation of pattern but not component tuned motion circuits in V1"

### Supplemental Figure 1

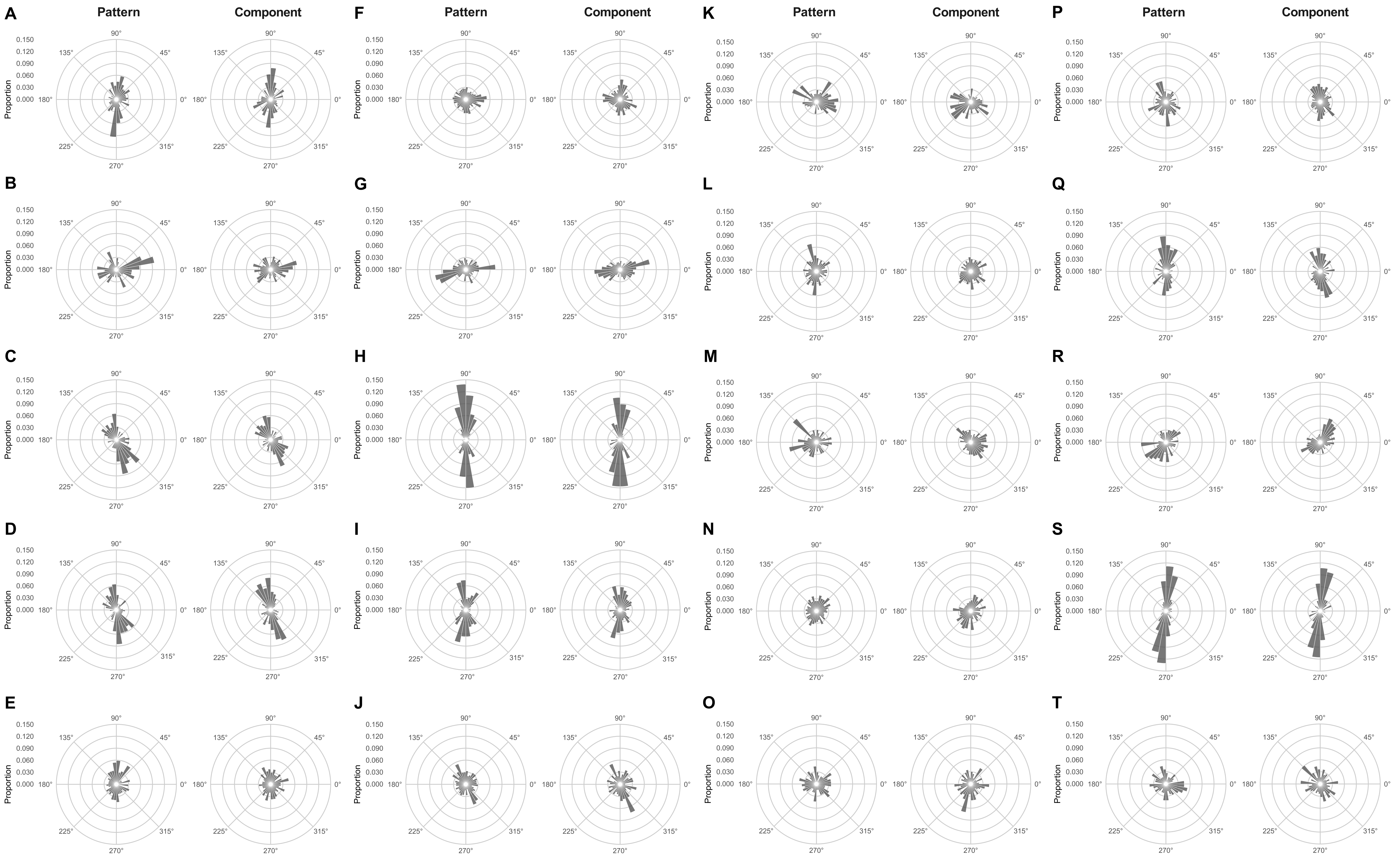
